# Zfp719 is a transcription factor important for maintenance of hearing in mice

**DOI:** 10.64898/2026.06.15.732256

**Authors:** Jing Chen, Neil J. Ingham, Maria Lachgar-Ruiz, Karim Boustani, Morag A. Lewis, Karen P. Steel

**Affiliations:** Wolfson Sensory, Pain, and Regeneration Centre, King’s College London, London, UK

## Abstract

Zfp719 is a zinc finger transcription factor which, when mutated, results in hearing impairment in mice. Its closest human orthologue, OTK18, has been linked to tinnitus in large human cohorts. Here we present our investigation of the electrophysiological, structural and transcriptional phenotypes in *Zfp719^tm1a^* mutant mice. Homozygotes have near-normal hearing at two weeks old, but lose sensitivity rapidly between two and three weeks, suggesting that while Zfp719 is not required for development, it is important for maintaining hearing. Heterozygous mice exhibit progressive hearing impairment for high frequencies at older ages. We observed damaged and degenerating outer hair cells from as early as three weeks old in homozygotes. We carried out bulk RNAseq at three ages and found one gene consistently misregulated, a long non-coding RNA specific to mice, *Gm15083*. A better understanding of the genes regulated by Zfp719 may shed light on genes and proteins important for maintaining hearing in humans.

## Introduction

The human gene *ZNF175* is a C2H2 zinc finger transcription factor, and encodes the protein OTK18. OTK18 has been proposed as an HIV type 1-inducible transcriptional suppresser as it is copiously expressed in macrophages following HIV type I infection [1], and may also play a role in innate immunity and in neurotrophin production, supporting neuronal survival [2]. More recently, mutations in *ZNF175* have been found to be associated with tinnitus in an analysis of several large human cohorts [3], making it a gene of interest in studies of the genetics of hearing impairment.

The C2H2 zinc finger genes (C2H2-ZNF) are one of the largest gene families in mammals, and are often found in clusters in the genome which show evidence of species-specific expansion and loss. In phylogenetic analyses of C2H2-ZNF genes, *ZNF175* belongs to a cluster of human genes which also includes *ZNF577*, *ZNF432*, *ZNF614*, *ZNF613*, *ZNF615*, *ZNF350*, and *ZNF649* [4]. The closest mouse orthologue to *ZNF175* is *Zfp719*, which belongs to a small rodent-specific subgroup of C2H2-ZNF genes closely linked to the *ZNF175* cluster [4]. Zfp719 and OTK18 share 44% amino acid identity.

Mice carrying a targeted mutation disrupting translation of *Zfp719* (*Zfp719^tm1a(EUCOMM)Wtsi^*, hereafter referred to as *Zfp719^tm1a^*) showed elevated auditory brainstem response (ABR) thresholds in a high-throughput targeted mutagenesis programme [5]. No other abnormalities were reported from this broad-based phenotyping pipeline (https://www.mousephenotype.org/data/genes/MGI:2444708). Here we present our study characterising the maturation and function of the inner ear of these mutant mice.

## Methods

### Ethical statement

Experiments involving mice were carried out in accordance with UK Home Office regulations and the UK Animals (Scientific Procedures) Act of 1986 (ASPA) under a UK Home Office licence, and the study was approved by the King’s College London Animal Welfare and Ethical Review Body. Mice were culled using methods approved under this licence to minimise any possibility of suffering. Mice were maintained in individually-ventilated cages at a standard temperature and humidity and in specific pathogen-free conditions, with lighting on a 12 hours on/12 hours of cycle, and in accordance with the EU Directive 2010/63/EU for animal experiments. Both sexes were used for this study. Experiments were carried out between 2 and 10 hours after lights on except for expression studies when samples were collected within a 1.5 hour window from 6 hours after lights on.

### Mice

The *Zfp719^tm1a(EUCOMM)Wtsi^* mouse line was generated at the Wellcome Sanger Institute as part of the Mouse Genetics Project, and was maintained on a C57BL/6N; C57BL/6N-A^tm1Brd^/a genetic background [6]. These mice are available from the European Mouse Mutant Archive (EM:06974). Animals were genotyped by PCR using template DNA extracted from pinna skin, by testing for the presence of the wildtype allele (forward primer: 5’-ATGGCTGTCACTGTCCCCTC-3’; reverse primer: 5’-TACCAGGAGGCCAGAAATGG-3’), a reaction which does not work in the mutant allele due to the size of the inserted cassette. The presence of the mutant allele was tested using the same forward primer with a reverse primer in the cassette (5’-TCGTGGTATCGTTATGCGCC-3’) and checked using primers against the neomycin resistance gene (forward primer: 5’-CAAGATGGATTGCACGCAGGTTCTC-3’; reverse primer: 5’-GACGAGATCCTCGCCGTCGGGCATGCGCGCC-3’) (Fig 1A).

**Figure 1.**
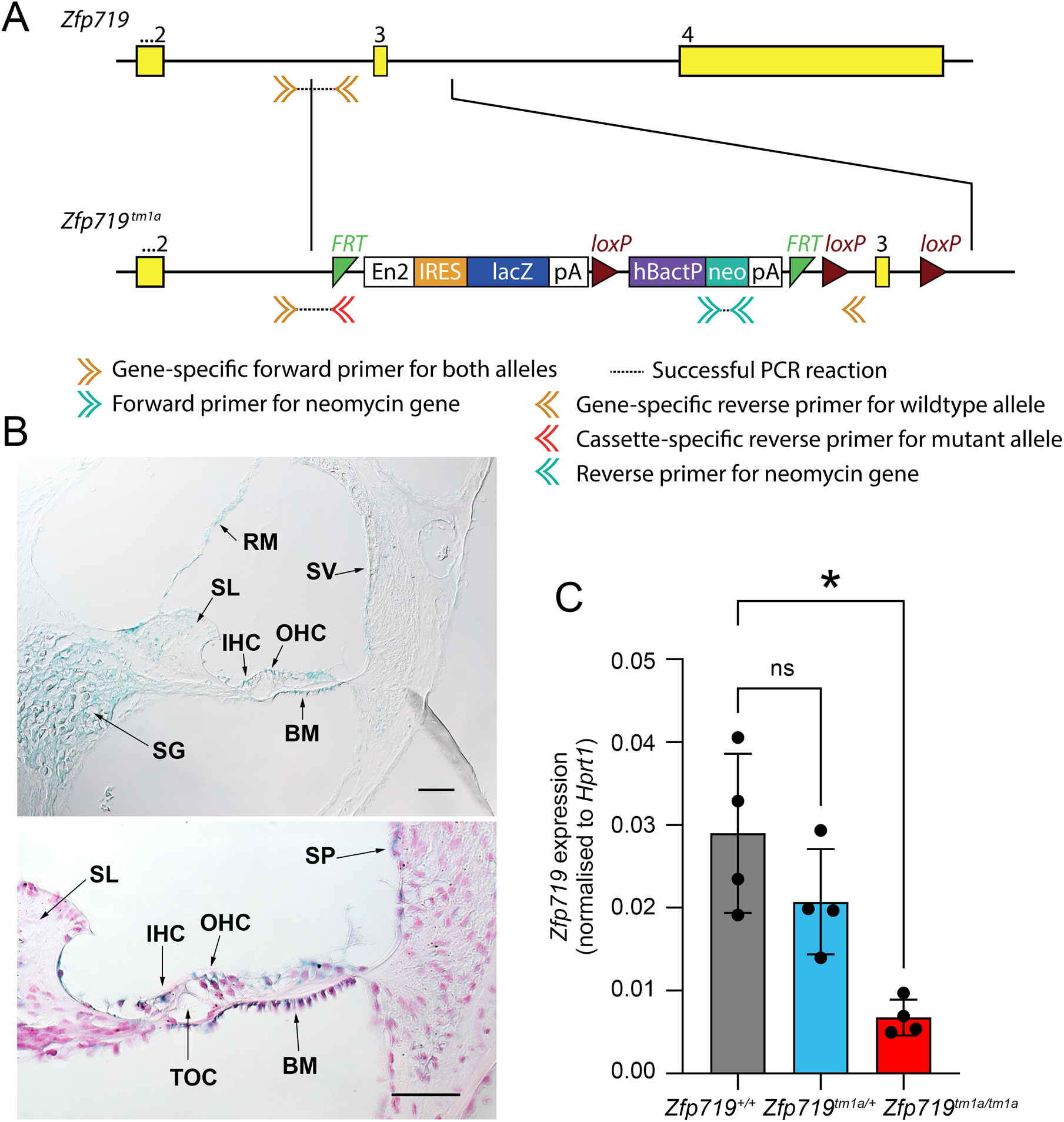
A) Diagram of the *Zfp719* tm1 a allele “knockout-first” design. The cassette, inserted between exons 2 and 3, interferes with normal transcription. Double arrowheads indicate the location and orientation of the primers used for genotyping. Yellow boxes: exons. Red triangles: loxP sites. Green triangles: FRT sites. Blue box: lacZ reporter gene. Teal box: neomycin resistance gene. Not to scale. B) X-gal staining (blue) at two weeks old in homozygotes (n=5; 3 heterozygotes and two wildtypes were also stained). The tissue was counterstained in pink (nuclear fast red, bottom panel). Expression is visible in the spiral ganglion (SG), inner hair cells (IHC), outer hair cells (OHC), and stria vascularis (SV). Also labelled are the tunnel of Corti (TOC), basilar membrane (BM), spiral limbus (SL) and Reissner’s membrane (RM). Scale bar= 50mm. C) The effect of the tm1a cassette on expression of *Zfp719* in the inner ear was tested using qPCR at P14. *Zfp719* expression was significantly reduced in homozygotes (p=0.0353, Dunnett’s T3 multiple comparisons test following one-way ANOVA), with a smaller, non-significant reduction in heterozygotes, both compared to wildtype littermates (n=4 for all genotypes).

### Quantitative real time PCR

The left inner ear was dissected from two week-old mice and stored in RNAlater (Life Technologies) at 4°C. RNA was extracted from inner ears using the QIAgen RNeasy Mini Kit (QIAgen, cat no 74104). cDNA was produced from normalised aliquots of RNA using SuperScript Reverse Transcriptase II (Invitrogen, cat no 18064014).

qPCR was performed using SsoAdvanced Master Mix (Applied Biosystems, cat no 1725284), and Taqman probes against *Zfp719* (cat no Mm01197328_m1) and *Hprt1* (cat no Mm01318747_g1), using a CFX Connect (BioRad). Relative gene expression was calculated using the 2^-ΔCT^ method [7].

### X-gal staining

The right inner ears from the same two week-old mice as above were collected and fixed in 4% paraformaldehyde (PFA) at 4°C for 45 minutes at room temperature, followed by PBS washes and decalcification in 0.1M EDTA overnight at 4°C. Samples were treated with X-gal detergent solution (2mM MgCl_2_, 0.02% NP-40; 0.01% sodium deoxycholate, 0.1M sodium phosphate buffer pH 7.3) for 30 minutes at room temperature (RT) with agitation, then incubated in staining solution (detergent solution plus 5mM K_3_Fe(CN)_6_, 5mM K_4_Fe(CN)_6_, and 1mg/ml X-gal in DMSO) at 37°C for 8 hours. After PBS washes, stained samples were embedded in paraffin wax and sectioned (8µm) onto Superfrost Plus slides. Chosen slides were dewaxed and mounted in DPX mounting medium. Some slides were counter-stained with nuclear Fast Red before mounting.

### Auditory Brainstem Response (ABR)

The hearing of wildtype and mutant *Zfp719^tm1a^* mice was tested using the ABR, as described in [8]. In brief, mice were sedated using a ketamine/xylazine mix (10mg ketamine and 0.1mg xylazine in 0.1ml per 10g body weight), and three subcutaneous needle electrodes were placed for recording the responses (reference over the left bulla, ground over the right bulla, and active electrode on the top of the head). For every stimulus (broadband click and 3kHz, 6kHz, 12kHz, 18kHz, 24kHz, 30kHz, 36kHz and 42kHz pure tone levels) and intensity from 0-95dB in 5dB steps, ABRs were recorded as the averaged evoked potential recorded from 256 stimulus presentations. Recovery from sedation was aided using atipamezole (0.01mg atipamezole in 0.1ml per 10g body weight). The threshold for each stimulus was defined as the lowest intensity at which a waveform could be identified. Mice were tested at 2, 3, 8, 14 and 26 weeks old; for the younger ages (2-4 weeks), ABR testing took place on the exact day (14 days, 21 days and 28 days post-natal). At 8 weeks, mice were tested at ages between 55 and 57 days; at 14 weeks, between 93 and 98 days; at 26 weeks, between 178 and 182 days. The cohort tested at 2 weeks was a separate cohort from the mice which underwent repeated ABR testing between 3 and 26 weeks.

Statistical tests were carried out using SPSS v31 (IBM). ABR thresholds were arcsine transformed and analysed using separate linear models for each stimulus frequency with a compound symmetric covariance structure and restricted Maximum Likelihood Estimation [9]. This method allows all data to be included, even when not all measurements are available for all subjects (for example, if a mouse dies before reaching 26 weeks) [10]. The interaction of genotype and age were considered for each stimulus, and data were corrected for multiple testing using the Bonferroni correction.

### Distortion Product Otoacoustic Emissions (DPOAEs)

DPOAEs were recorded from anaesthetised mice as described in [11]. Briefly, stimulus tones of 6, 12, 18, 24 or 30kHz (f2) and corresponding f1 tones (in a ratio of 1:1.2 for f1:f2) were presented into the mouse ear canal. Stimuli were presented over f2 levels from -10dB to 65dB sound pressure level in 5dB steps, with f1 set to 10dB higher than f2. The sound present in the mouse ear canal was recorded using an Etymotic ER10B+ low noise microphone, and the resulting waveform was subjected to Fast Fourier Transform to extract the amplitude of the 2f1-f2 DPOAE component and the microphone noise floor for each frequency. The 2f1-f2 component was plotted against the f2 stimulus levels, and the DPOAE threshold was estimated as the lowest f2 stimulus level where the DPOAE exceeded 2 standard deviations above the mean noise floor sound pressure level. Statistical tests were carried out using the same linear mixed model method as for the ABR data.

### Endocochlear potential (EP) measurements

Mice were anaesthetised with urethane (0.1ml / 10g of a 20% solution, via ip injection), a tracheal cannula inserted and the left bulla opened to expose the wall of the cochlea. A small hole was made over the basal turn of the cochlea and the tip of a 150mM KCl-filled glass microelectrode was inserted into the scala media. The endocochlear potential was measured as a stable positive potential, referenced to a Ag-AgCl pellet placed under the dorsal skin of the neck [12]. Once the stable resting (normoxic) EP had been recorded, the electrode was left *in situ* and the mouse injected with an overdose of urethane. Following onset of anoxia, the potential fell rapidly and the maximally negative (anoxic) EP was measured [12]. A one-way ANOVA with Tukey’s multiple comparison test was used for statistical analysis, carried out using GraphPad Prism.

### Scanning electron microscopy (SEM)

Inner ears were dissected from mice aged either 3 or 8 weeks, and placed in fixative solution (0.1M sodium cacodylate; 3uM calcium chloride; 2.5% glutaraldehyde) for 3 hours at RT. Fixed samples were washed with PBS before being dissected to expose the organ of Corti by removing the surrounding bone, lateral wall and Reissner’s membrane. Samples were then post-fixed with osmium tetroxide (1%) then thiocarbohydrazide, repeated three times (OTOTO technique; [13]) followed by dehydration in graded ethanol (20% - 100%) and critical point drying. Images were taken of the samples using a FEI Quanta 200F scanning electron microscope or a JEOL JSM 7800 Prime scanning electron microscope. For counting, three images were taken at the 12kHz and 30kHz best frequency regions to capture approximately 90 outer hair cell bundles at each location. Hair bundles were classified as “unaffected”, “lost” or “fusion”, depending on whether they appeared normal, were absent, or exhibited fused stereocilia. One ear was used from each mouse.

### Wholemount dissection, immunohistochemistry, and confocal imaging of presynaptic puncta

Cochleae of 3 week-old mice were fixed in 4% paraformaldehyde (PFA) in PBS for 2h, decalcified with 0.1M ethylenediaminetetraacetic acid (EDTA) overnight at RT, samples were permeabilized with 5% Tween in PBS for 30 min and blocked in 0.5% Triton X-100 and 5% Horse Serum (HS) in PBS for 1h.

Primary antibody staining took place overnight at 4°C, with rabbit anti-Myosin VIIa (diluted 1:200; 25-6790, Proteus) and mouse IgG1 anti-C-terminal binding protein 2 (CtBP2) (diluted 1:200; 612044, BD Transduction Laboratories), diluted in 0.5% Triton X-100 and 1% HS in PBS. Secondary antibody staining was carried out for 1h at RT, with Alexa Fluor 647-conjugated chicken anti-rabbit (1:200; #A21443, Thermo Fisher Scientific) and Alexa Fluor 568-conjugated goat anti-mouse (IgG1) (1:1000; #A21124, Thermo Fisher Scientific). Samples were mounted using ProLong Gold Antifade Mountant with DAPI (P36931, Life Technologies) and imaged using a Zeiss Imager 710 confocal microscope connected to ZEN 2010 software. All images were captured using a plan-Apochromat 63x/1.40 oil DIC objective with a 2.0 optical zoom. Confocal z-stacks were acquired with a z-step size of 0.25 μm to ensure imaging of all synaptic puncta. Images were acquired at the 12 kHz and 30 kHz best-frequency regions. The ImageJ measure line plug-in from Eaton Peabody Laboratories was used to map cochlear length to cochlear best frequencies. Brightness and contrast levels were adjusted for whole images using Fiji software.

Manual counting of synaptic puncta was carried out using the cell-counter plugin in Fiji software. All images containing puncta were merged into a z-stack, and the z-axis maximum intensity projection was used to quantify presynaptic puncta. The number of ribbon synapses was divided by the number of Myo7a-labelled IHCs in the image to determine the number of ribbons per IHC. In cases where an IHC was not completely visible in the image, the corresponding puncta were not included in the count. Two non-overlapping images were acquired for each best-frequency region, with a count of 14 to 20 IHCs at each frequency. The three genotype groups were compared using a one-way ANOVA with Tukey’s multiple comparison test, carried out using GraphPad Prism.

### Sample preparation for RNAseq

Mice from heterozygote x heterozygote matings were collected at 4 days, 2 weeks and 3 weeks old and culled by decapitation (for 4 day-old mice) or cervical dislocation (for 2 and 3 week-old mice). The head was bisected and the brain removed. For 4 day-old mice, the inner ears were dissected out in RNAlater. For the older ages, the inner ears were isolated from the temporal bones before moving into RNAlater, then followed by a further dissection in RNAlater to get rid of extra tissue attached to the inner ears. The inner ears were transferred into eppendorf tubes (without RNAlater) and flash-frozen in liquid nitrogen. After genotyping using pinna tissue DNA, the inner ears from sex matched littermates were used. RNA was extracted using the QIAgen RNeasy kit with QIAshredder columns (cat no 79654).

### RNAseq

Strand-specific libraries were prepared using the Illumina TruSeq stranded mRNA LT kit. Sequencing was carried out on an Illumina NextSeq 500 machine as paired-end 150bp reads. Reads were filtered using filterbytile (from BBMap_37.33) [14] and trimmed with Trimmomatic 0.36 [15]; adapters were removed, trailing ends were clipped where the quality was low and a sliding window approach used to control for quality across the entire read. Reads with 36 basepairs or fewer were discarded. Reads were assembled to GRCm38 using Hisat2 version 2.0.2beta [16]. Bam files were soft-clipped beyond end-of-reference alignments and MAPQ scores set to 0 for unmapped reads using Picard 2.10.0. The count matrixes were generated by htseq v2.0.3 [17]. The edgeR package [18] was used to perform a generalised linear model likelihood ratio test. The resulting gene lists, including differential expression between homozygotes and wildtype samples and false discovery rate, are shown in Supplementary Table S1.

### Protein and gene sequence cross-species analyses

Protein (Zfp719; ENSMUSP00000050968.8) and DNA (*Gm15083*; ENSMUST00000138670.3) sequences were obtained from the Ensembl database (release 115) [19]. Ensembl’s BLASTN and BLASTP tools were used to search DNA and protein sequences respectively against a list of genomes chosen from those available (see Supplementary Table S2 for all species). For both sequences, a good match was defined as a high score (>1000) and full or nearly full-length alignment (735 amino acids for the Zfp719 protein, 1413 base pairs for the *Gm15083* transcript). The first exon of *Gm15083* (ENSMUSE00000754815) was also used in a separate BLASTN search against all Muroidea genomes, because it was too short to be found separately in the *Mus caroli* genome using the full-length sequence. While a match for this exon was confirmed in the *Mus caroli* genome, it was not found in any other Muroidea which did not have a full-length match.

## Results

### *Zfp719* is expressed in multiple locations in the inner ear

The *Zfp719^tm1a^* allele contains a large cassette inserted in the intron between exons 2 and 3, which disrupts transcription leading to a knockdown [20] (Fig 1A). The cassette includes the LacZ reporter which can be used for expression analysis using X-gal staining. *Zfp719* was found to be expressed throughout the inner ear at post-natal day (P)14, including the hair cells, stria vascularis, and spiral ganglion neurons (Fig 1B). qPCR analysis showed that expression is reduced (to around 23% of normal levels) in homozygotes (Fig 1C), although it is not eliminated, suggesting that some transcription of the full-length mRNA still takes place despite the presence of the cassette. This “leaky” expression has been reported from other knockout-first alleles [6, 21, 22]. There was an intermediate level of expression in heterozygous mice compared with wildtypes (Fig 1C).

### *Zfp719* homozygotes exhibit rapid progressive hearing loss

Mice carrying the mutant allele showed close to normal ABR thresholds at two weeks old, but lost hearing sensitivity rapidly between two and three weeks. By 6 months old homozygotes were completely unresponsive to sounds, while heterozygotes showed high frequency hearing impairment (Fig 2A). The function of outer hair cells was assessed using DPOAEs; impaired outer hair cell (OHC) function was observed at 4 weeks old but not at 2 weeks, correlating with the progressive change in ABR thresholds (Fig 2B).

**Figure 2.**
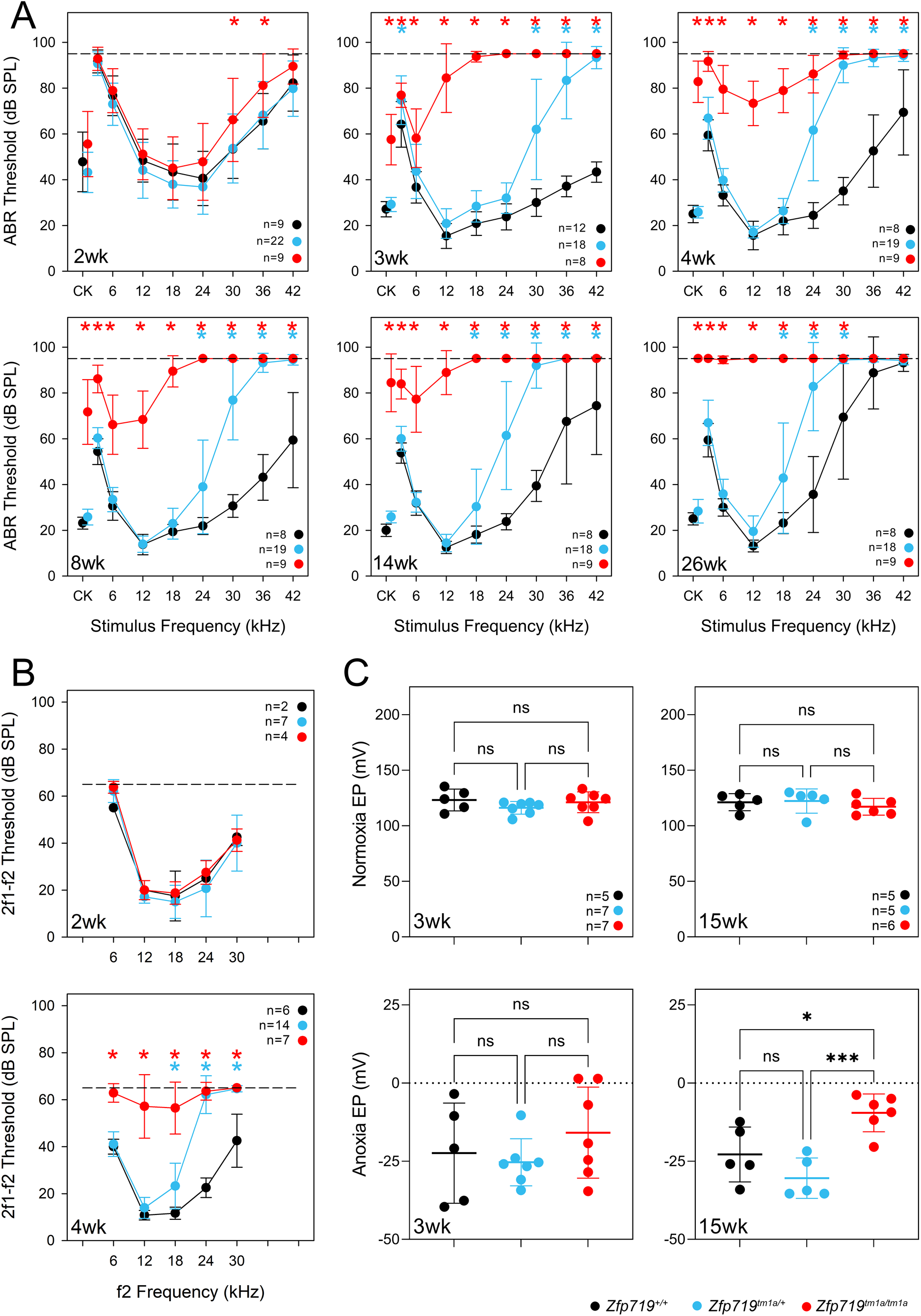
A) Average ABR thresholds for *Zfp71*_gtmia_ wildtype (black), heterozygous (blue) and homozygous (red) mice at ages between `2` and `26` weeks. Homozygotes lost sensitivity rapidly from 2 to 3 weeks old and hearing deteriorated with age until there was no response to any frequency tested at 26 weeks old (6 months). Heterozygotes showed high frequency hearing loss at 3 weeks old, but hearing loss progressed less rapidly than in homozygous littermates. B) DPOAEs were normal at 2 weeks old but abnormal at 4 weeks old. For A and B, asterisks indicate statistically significant differences between heterozygotes and wildtypes (blue) or homozygotes and wildtypes (red) (Bonferroni-corrected p < 0.05, mixed linear model pairwise comparison). C) Normal endocochlear potentials (EP) were recorded at 3 weeks and 15 weeks old in mice of each genotype, indicating normal strial function. Anoxic endocochlear potentials showed a less negative EP in the homozygous mutants at 15 weeks old. At least 5 animals were tested at each age and for each genotype. *p=0.022; ***p=0.0009 (one-way ANOVA, followed by Tukey’s multiple comparisons test).

Normoxic and anoxic endocochlear potential measurements were conducted at 3 weeks and 15 weeks old. There was no significant difference among homozygous, heterozygous and wildtype mice at 3 weeks old in either measure, although the minimum anoxic potential was relatively variable in all genotypes. At 15 weeks old, the normoxic endocochlear potential was unaffected, but the anoxic EP was significantly less negative in the homozygotes (Fig 2C).

### Outer hair cells are severely affected in homozygotes

Scanning electron microscopy on mice at three weeks old revealed that hair cells of heterozygotes appeared normal at both the 12kHz and 30kHz best frequency regions. In homozygotes at three weeks old, OHCs appeared normal at the 12kHz region but abnormalities such as missing or fused stereocilia were evident at the 30kHz region, and a small number of OHCs were missing (Fig 3A). However, by 8 weeks old, fused stereocilia were also visible in heterozygotes (Fig 3B), while extensive hair cell degeneration was observed in homozygotes (Fig 3A, B).

**Figure 3.**
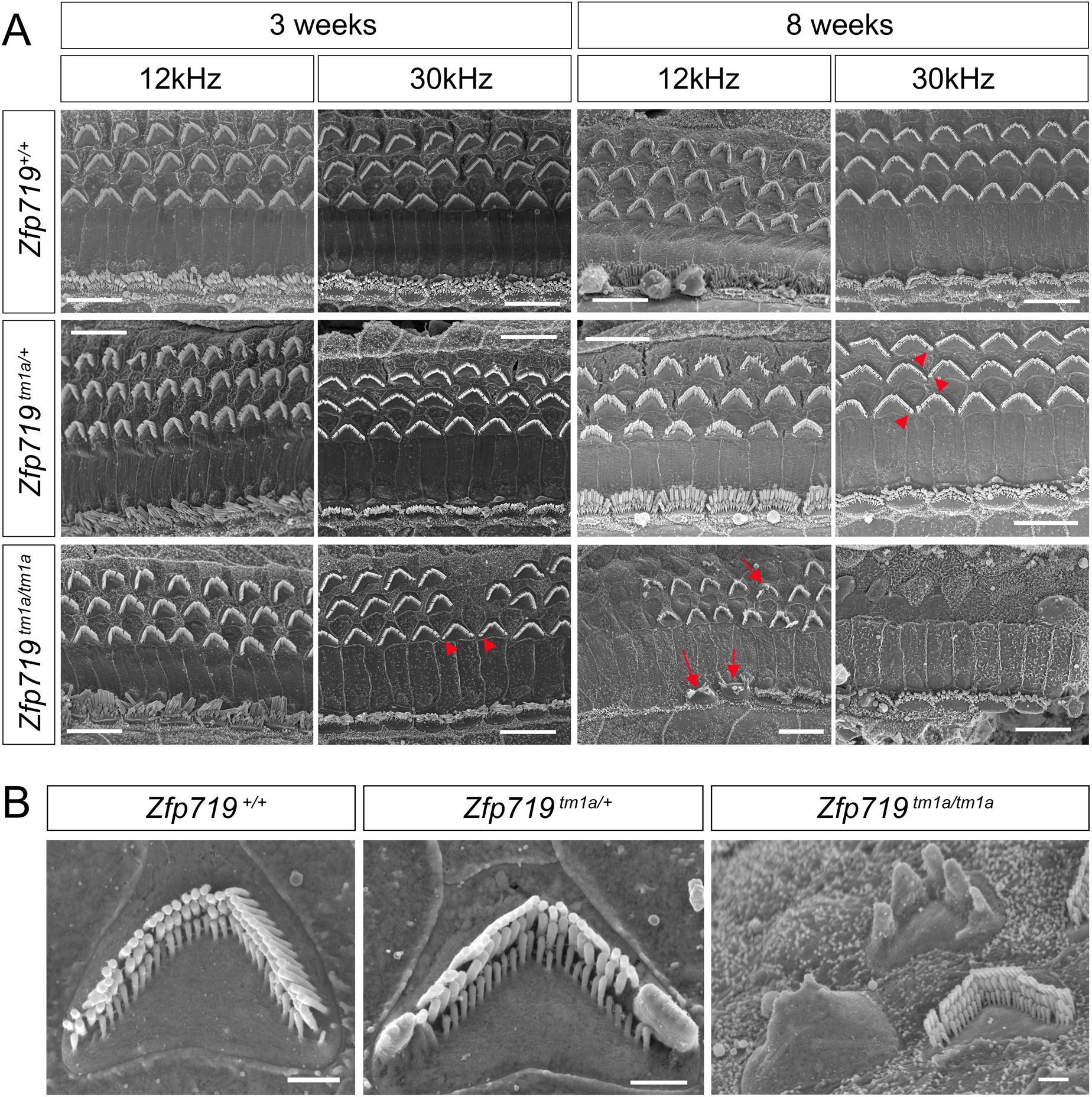
A) Scanning electron micrographs of the organ of Corti of wildtype, heterozygous and homozygous mice at 3 weeks and 8 weeks. Hair cells were present in homozygotes at 3 weeks (n=5), but OHCs showed subtle defects including fused stereocilia (red arrowheads), and some were missing their stereocilia bundles. IHCs appeared normal at this age. Heterozygotes (n=9) looked similar to wildtypes (n=3). At 8 weeks, homozygotes (n=4) showed more severely fused stereocilia bundles of both IHCs and OHCs (arrows), more missing bundles, and patches where all stereocilia bundles were missing, which was more common towards the base. Fused stereocilia were visible in heterozygotes (n=3) at this stage (red arrowheads), but no abnormalities were seen in wildtypes (n=2). Scale bar= 10µm. B) Close-ups of outer hair cells from wildtype, heterozygous and homozygous mice at 8 weeks old, showing stereocilia fusion in the heterozygote and different stages of degeneration in the homozygote. Scale bar= 1µm.

### There are no significant differences in synaptic ribbon counts at three weeks old

As the ribbon synapses of inner hair cells can be reduced in some situations, we carried out immunofluorescent labelling of the organ of Corti at three weeks old, an age when ABR thresholds are raised in homozygous *Zfp719^tm1a^* mutants. Hair cells were labelled with Myo7a, and presynaptic ribbons with CtBP2 antibodies. We counted the number of labelled puncta per inner hair cell at the 12kHz and 30kHz best-frequency regions and found no significant differences between the genotypes in either region (Fig 4).

**Figure 4.**
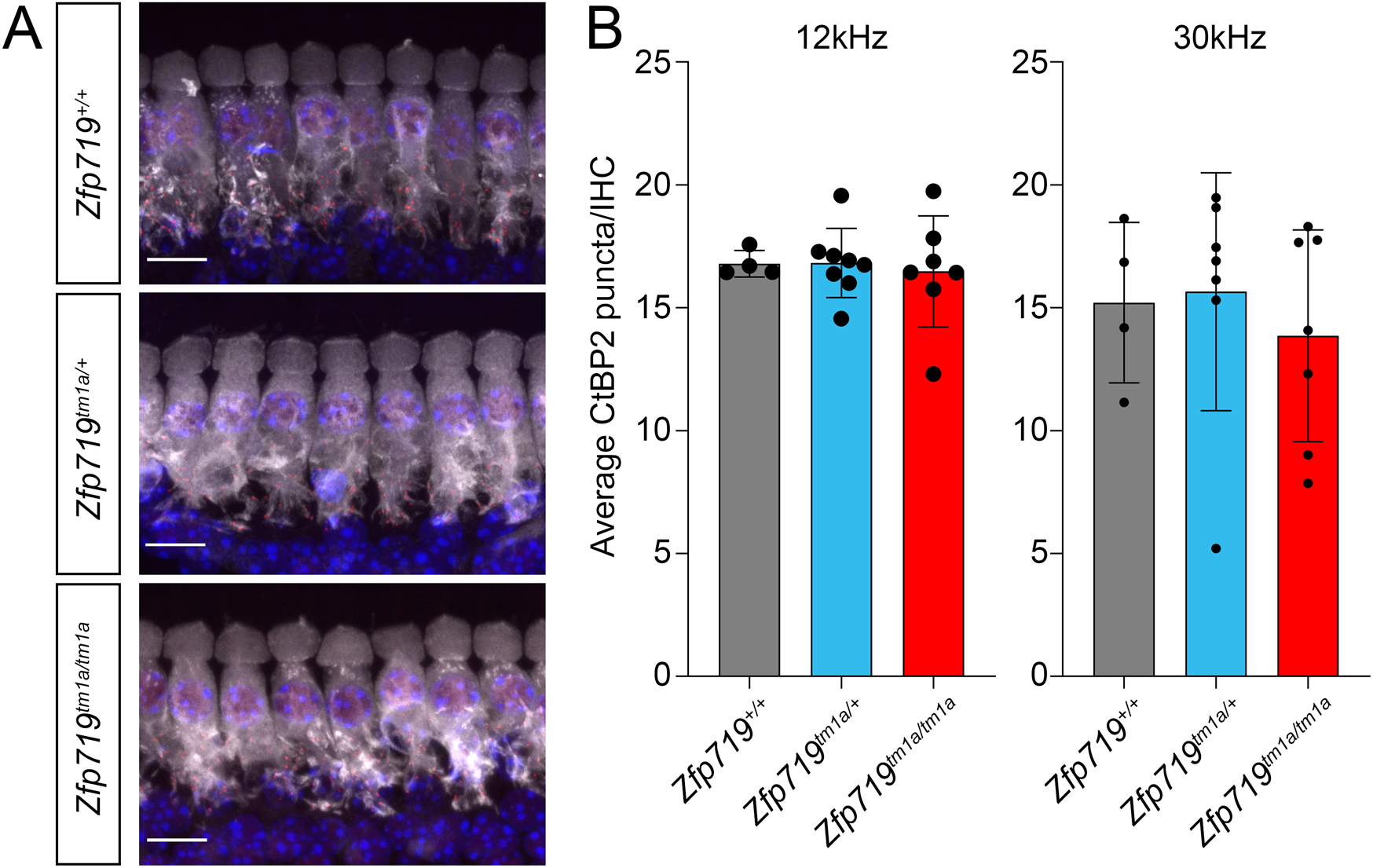
A) Confocal images showing IHCs (Myo7a; white) and presynaptic ribbons (Ctbp2; red) in a wildtype, a heterozygote, and a homozygote at three weeks old. Nuclei are stained with DAPI (blue). Scale bar= 10 µm. B) Bar chart with individual data points showing the average count of CtBP2 puncta per inner hair cell at the 12kHz and the 30kHz region in wildtypes (n=4), heterozygotes (n=8 at 12kHz, n=7 at 30kHz) and homozygotes at three weeks old (n=7).

### Not many genes are misregulated in *Zfp719^tm1a^* homozygotes

Zfp719 is a transcription factor, so in order to investigate the genes it regulates, we carried out RNA-seq using RNA extracted from inner ear samples of wildtype, heterozygous and homozygous littermates at 4 days old, two weeks old and three weeks old. We found eight significantly misregulated genes at 4 days old (the developmental stage) and three significantly misregulated genes at two weeks old (before onset of hearing loss), while at three weeks old (after the onset of hearing loss), there were six misregulated genes. The only genes common to all three ages were *Zfp719* itself which was downregulated in homozygous mutants, and a long non-coding RNA gene called *Gm15083* which was upregulated in mutant inner ears (Fig 5).

**Figure 5.**
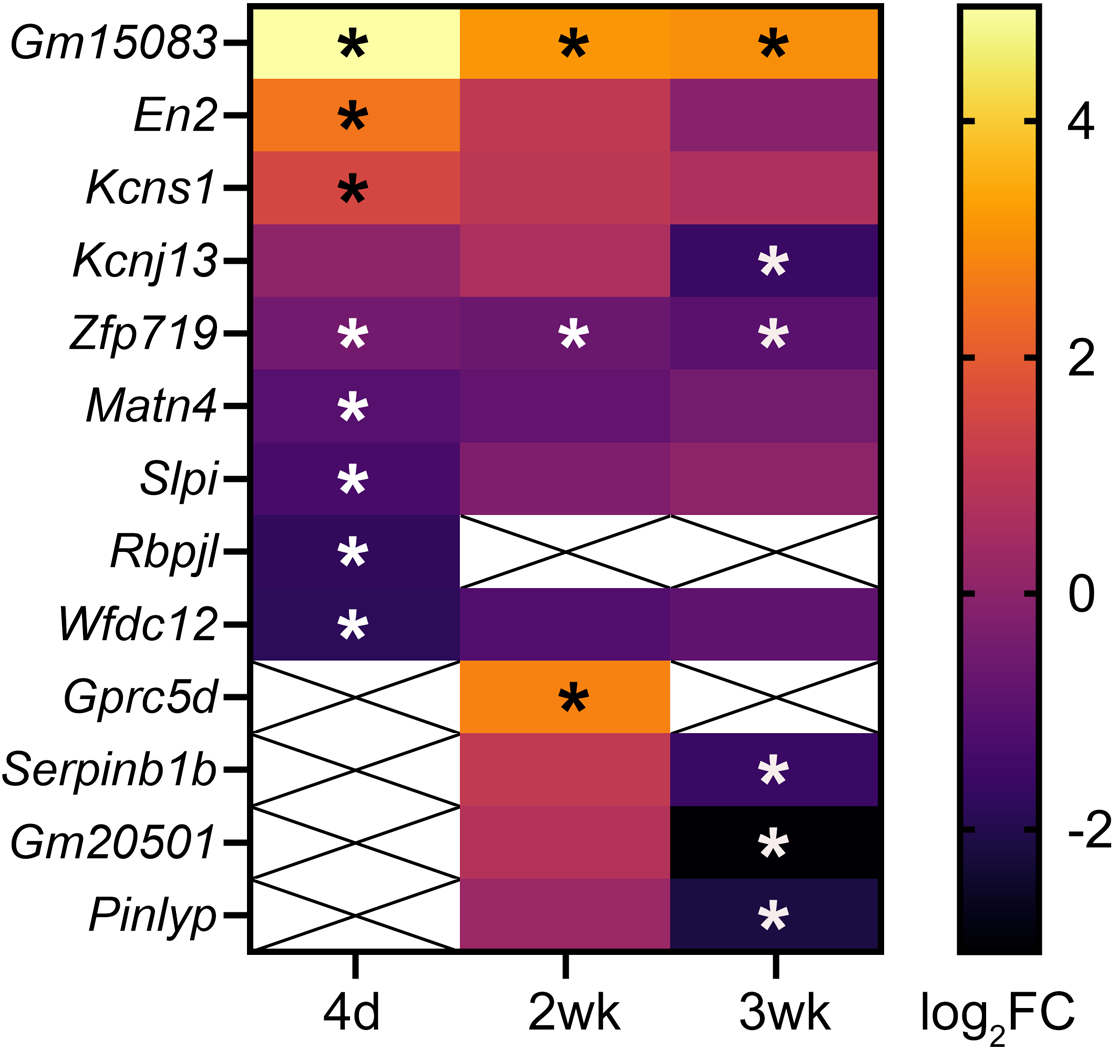
Heatmap showing all the genes which were significantly misregulated at one or more ages. The colour scale shows the log2 of the fold change in homozygotes compared to wildtypes. Asterisks indicate significant misregulation (FDR< 0.05). Some genes were not reliably detected at some ages, and have not been included; indicated by crossed-out white boxes (eg *Gprc5d* at P4 and P21).

### Zfp719 and *Gm15083* are phylogenetically restricted

*Gm15083* is a predicted long noncoding RNA gene with only two known orthologues (Ensembl database, accessed May 2026), from the shrew mouse (*Mus pahari*) and the western wild mouse (*Mus spretus*). We used Ensembl BLAST with the sequence of the mouse reference for the *Gm15083* transcript (ENSMUST00000138670.3) and found hits for all *Mus musculus* substrains, including *Mus musculus domesticus*, *Mus musculus castaneous*, and *Mus musculus molossinus*. We also found matches in the Steppe mouse (*Mus spicilegus*) and Ryukyu mouse (*Mus caroli*) genomes, although they were not annotated as genes in Ensembl. We did not find high quality matches against any other genomes, including the rat and other rodents (Supplementary Table S2).

We then investigated matches to the protein sequence of *Mus musculus* Zfp719 (ENSMUSP00000050968.8), using Ensembl BLASTP. We found matches in a wider range of rodents, including rat (*Rattus norvegicus*), golden hamster (*Mesocricetus auratus*) and prairie vole (*Microtus ochrogaster*), but nothing in other species, including less related rodents such as the guinea pig (*Cavia porcellus*) and the degu (*Octodon degus*), and a variety of other eukaryotic genomes, including human, zebrafish, fruit fly and yeast (Supplementary Table S2). We plotted the presence of these sequences against the Ensembl Species Tree (http://sep2025.archive.ensembl.org/info/about/speciestree.html) and found that while *Zfp719* appears to have arisen in the Muroidea clade (Taxon ID 337687), *Gm15083* was found only in the Mus subgenus (Taxon ID 862507) (Fig 6).

**Figure 6.**
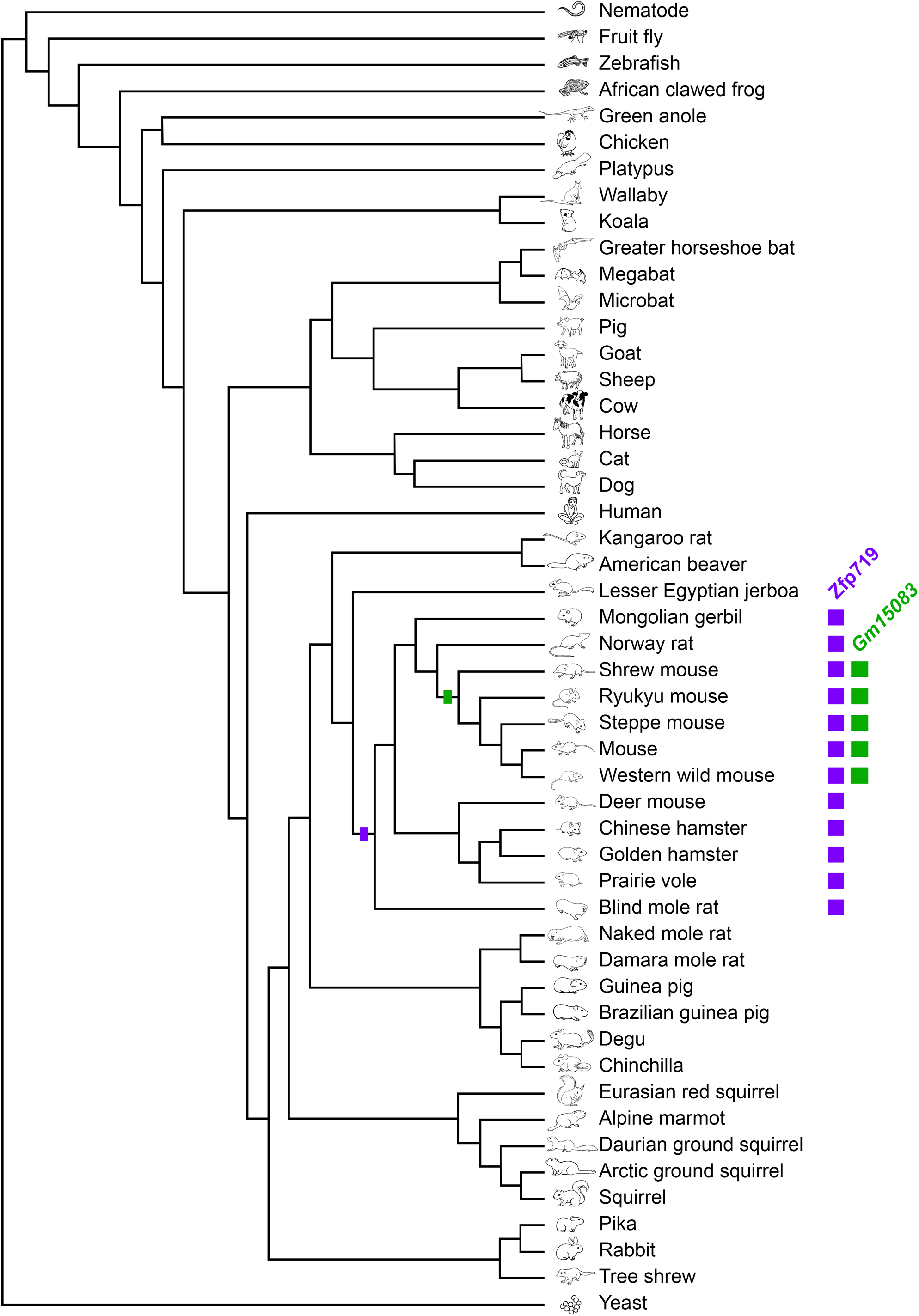
Adaptation of the Ensembl Species Tree (http://sep2025.archive.ensembl.org/info/about/speciestree.html) showing just those genomes which were tested for the presence of Zfp719 (ENSMUSP00000050968.8) and *Gm15083* (ENSMUST00000138670.3). The purple bar indicates where Zfp719 arose, and the green bar where *Gm15083* arose. Purple and green to the right mark where each is present in a species genome.

## Discussion

Zfp719 was found to be expressed throughout the mouse cochlea, but the first morphological change in these mutants appears to be in outer hair cells, followed later by inner hair cells. However, at three weeks old, when thresholds were highly raised in homozygotes (Fig 2A), OHC stereocilia showed only subtle morphological defects (Fig 3), and the ribbons of IHC synapses were present in normal numbers, suggesting that the underlying defect causing raised ABR thresholds at three weeks old remains to be fully understood.

Although Zfp719 expression is seen in the stria vascularis, we did not observe any evidence of effects on the normoxic endocochlear potential at 3 or 15 weeks old in our study (Fig 2C). We did observe a difference in the anoxic EP seen in 15 week old homozygotes (Fig 2C). The negative endocochlear potential likely results primarily from the rapid leakage of cations from the scala media through spontaneous openings of the mechanoelectrical transduction channels in the IHCs and OHCs. A less negative potential suggests there is a reduction in the rate of cation leakage out of scala media. Given that hair cell loss is obvious from 8 weeks (Fig 3), we suggest that this alteration in anoxic EP is likely due to the loss of hair cells and the concomitant loss of mechanoelectrical transduction channels.

Although Zfp719 is a transcription factor, we saw a limited effect on the transcriptome at the three ages we studied (Fig 5). We carried out bulk RNAseq on the whole inner ear, and it may be that single cell RNAseq will allow more precise assessments of the effects of mutations in *Zfp719* in specific cell types. Only one of the significantly misregulated genes has been previously associated with hearing impairment; *Kcnj13*, which is downregulated in *Zfp719^tm1a^* homozygotes at three weeks old. Mice carrying a *Kcnj13* conditional knockout allele have been described as showing severe hearing loss as adults (Xu et al 2022) [23]. This paucity of links to existing deafness-associated genes and regulatory networks offers few clues as to the underlying mechanisms of *Zfp719*-associated hearing impairment. The one gene which was consistently upregulated in all three ages, *Gm15083*, is a predicted gene whose only orthologues were found in other species of mouse (Fig 6). If *Gm15083* is involved in mediating the effect of Zfp719 in the inner ear, it may be a mouse-specific function, but the presence of Zfp719 in a wider range of rodents (Fig 6) suggests it regulates other genes besides *Gm15083*.

Mice are commonly used in hearing research due to the similarity of their ears to human ears, structurally, functionally, and genetically, in development and in maintenance of hearing. Many genes known to be involved in human hearing impairment also underlie hearing loss in mice [24]. In this case, the absence of direct Zfp719 orthologues outside the Muroidea clade might suggest the *Zfp719^tm1a^* mutant mouse is not a good model for studying human hearing. However, even if the role of Zfp719 in the inner ear is specific to rodents, or even specific to mice, it is likely that the genes it regulates connect to conserved regulatory networks further downstream which are relevant to human hearing. This is supported by the link between tinnitus and OTK18 [3], the closest human orthologue to Zfp719. Further research into the factors controlled by Zfp719 in the mouse inner ear would help elucidate this.

The development of normal hearing at an early age prior to rapid progressive hearing loss suggests that Zfp719 is not critical for development of the inner ear and auditory function, but is necessary for maintaining hearing. Even the comparatively small difference in *Zfp719* expression seen in the heterozygotes (Fig 1C) is sufficient to cause later-onset progressive hearing impairment (Fig 2A), suggesting the role that Zfp719 plays is an important one. Even if the factors controlled by Zfp719 in the mouse inner ear do not directly map to orthologues in the human inner ear, understanding the underlying systems may shed light on other proteins required for maintaining hearing in ageing, in humans as well as in mice.

## Author contributions

Conceptualisation: KPS; Data curation: JC, NJI, MAL; Formal analysis: JC, NJI, MLR, KB, MAL; Funding acquisition: KPS, NJI, MAL; Investigation: JC, NJI, MLR, KB, MAL; Methodology: NJI, MAL; Project administration: KPS; Supervision: KPS; Writing – original draft: JC, MAL; Writing – review and editing: JC, NJI, MLR, KB, MAL, KPS.

## Acknowledgements

We thank the King’s College London Centre for Ultrastructural Imaging for access to the scanning electron microscopy facilities. We thank the Wellcome Sanger Institute Mouse Genetics Project for generating and providing the *Zfp719^tm1a^* mutant mice.

## Data availability

The RNAseq data are available from ArrayExpress, accession E-MTAB-7414.

---

**Supplementary Table S1.** Bulk RNA-seq expression data comparing homozygotes to wildtypes at P4, P14 and P21.

**Supplementary Table S2.** List of species tested for the presence of Zfp719 protein sequence (ENSMUSP00000050968.8) and *Gm15083* transcript sequence (ENSMUST00000138670.3) using Ensembl BLASTP and BLASTN respectively. Two species had matches for the *Gm15083* sequence which were not annotated as genes (*Mus caroli* and *Mus spicilegus*), and the genomic coordinates are provided for these instead of the gene ID.

